# Endothelial JAK2V617F mutation leads to thrombosis, vasculopathy, and cardiomyopathy in a murine model of myeloproliferative neoplasm

**DOI:** 10.1101/2019.12.31.891721

**Authors:** Melissa Castiglione, Christopher Mazzeo, Ya-Ping Jiang, Juei-Suei Chen, Kenneth Kaushansky, Wei Yin, Richard Z. Lin, Haoyi Zheng, Huichun Zhan

## Abstract

**Rational:** The myeloproliferative neoplasms (MPNs) are clonal hematological malignancies characterized by hematopoietic stem cell expansion and overproduction of mature blood cells. Cardiovascular complications are the leading cause of morbidity and mortality in patients with MPNs. The acquired kinase mutation JAK2V617F plays a central role in these disorders. Mechanisms responsible for cardiovascular dysfunction in MPNs are not fully understood, limiting the effectiveness of current treatment.

**Objective:** Vascular endothelial cells (ECs) play critical roles in the regulation of hemostasis and thrombosis. ECs carrying the JAK2V617F mutation can be detected in patients with MPNs. The goal of this study was to test the hypothesis that the JAK2V617F mutation alters endothelial function to promote cardiovascular complications in patients with MPNs.

**Methods and Results:** We employed murine models of MPN in which the JAK2V617F mutation is expressed in specific cell lineages. When JAK2V617F is expressed in both blood cells and vascular ECs, the mice developed MPN and spontaneous, age-related dilated cardiomyopathy with an increased risk of sudden death as well as a prothrombotic and vasculopathy phenotype on histology evaluation. We showed that JAK2V617F-mutant ECs are required for this cardiovascular disease phenotype and the mutation can alter endothelial cell function. Finally, in a more therapeutically oriented approach, we demonstrated that transplantation with wild-type donor marrow cells can improve cardiac function by reversing the left ventricle remodeling process in this JAK2V617F-positive MPN murine model.

**Conclusions:** These findings suggest that the JAK2V617F mutation alters vascular endothelial function to promote cardiovascular complications in MPNs. Therefore, targeting the MPN vasculature represents a promising new therapeutic strategy for patients with MPNs.

## INTRODUCTION

The chronic Philadelphia chromosome (Ph^1^) negative myeloproliferative neoplasms (MPNs), including polycythemia vera (PV), essential thrombocythemia (ET) and primary myelofibrosis (PMF), are clonal stem cell disorders characterized by hematopoietic stem/progenitor cell (HSPC) expansion and overproduction of mature blood cells. Patients with MPNs suffer from many debilitating complications including both venous and arterial thrombosis, with cardiovascular events being the leading cause of morbidity and mortality in these patients.^1^ The acquired kinase mutation JAK2V617F plays a central role in these disorders. Mechanisms responsible for cardiovascular diseases in MPNs remains incomplete, limiting the effectiveness of current treatment.

Recently, several studies have reported that JAK2V617F is one of the common mutations associated with clonal hematopoiesis of indeterminate potential (CHIP), which is present in more than 10% of people older than 70 yrs.^2–6^ Not surprisingly, those who harbor these clonal hematopoiesis have a modestly increased risk of developing hematologic cancers, such as acute leukemia. Unexpectedly, as few as 2% mutation-bearing blood cells can produce a 2-4 fold increase in heart diseases in individuals with CHIP, even in the absence of other risk factors for heart diseases. In particular, individuals with JAK2V617F-mutant CHIP have 12 times the risk of coronary heart disease and ischemic stroke compared to individuals without any CHIP-associated mutation.^4, 6^ Therefore, since up to 40% of patients with MPNs develop thrombosis,^1^ and MPN-associated mutation in seemingly healthy individuals is associated with an excess of cardiovascular morbidity and mortality, MPN provides an ideal model system to investigate the relationship between hematopoietic mutations and cardiovascular disorders.

Vascular endothelial cells (ECs) play critical roles in the regulation of hemostasis and thrombosis.^7^ ECs carrying the JAK2V617F mutation can be detected in isolated (by laser microdissection) liver or spleen ECs in 67% of patients with MPNs.^8, 9^ One study demonstrated that both wild-type and JAK2V617F-mutant spleen ECs co-exist in the same patients and mutant ECs comprise 40-100% of all splenic ECs.^9^ The mutation can also be detected in 60-80% of EC progenitors derived from the hematopoietic lineage and, in some reports based on *in vitro* culture assays, in endothelial colony-forming cells from MPN patients.^9–13^ Although vascular ECs in patients with MPNs are likely heterogeneous with both normal and JAK2V617F-mutant ECs, the available data is clear that mutant ECs are present in many or most patients with MPNs, albeit at variable frequency. In this work, we tested the hypothesis that the JAK2V617F mutation alters endothelial function to promote cardiovascular complications in patients with MPNs.

## MATERIALS AND METHODS

### Experimental mice

JAK2V617F Flip-Flop (FF1) mice^14^ was provided by Radek Skoda (University Hospital, Basal, Switzerland) and *Tie2-Cre* mice^15^ by Mark Ginsberg (University of California, San Diego). FF1 mice were crossed with Tie2-Cre mice to express JAK2V617F specifically in hematopoietic cells and ECs (Tie2^+/−^FF1^+/−^, or Tie2FF1 mice), so as to model the human diseases in which both the hematopoietic stem cells and ECs harbor the mutation. All mice used were crossed onto a C57BL/6 background and bred in a pathogen-free mouse facility at Stony Brook University. CD45.1+ congenic mice (SJL) were purchased from Taconic Inc. (Albany, NY). All mice were fed a standard chow diet. No randomization or blinding was used to allocate experimental groups. Animal experiments were performed in accordance with the guidelines provided by the Institutional Animal Care and Use Committee.

### Complete blood counts

Peripheral blood was obtained from the facial vein via submandibular bleeding, collected in an EDTA tube, and analyzed using a Hemavet 950FS (Drew Scientific).

### Transthoracic echocardiography

Transthoracic echocardiography was performed on mildly anesthetized spontaneously breathing mice (sedated by inhalation of 1% isoflurane, 1 L/min oxygen), using a Vevo 3100 high-resolution imaging system (VisualSonics Inc., Toronto, Canada). Both parasternal long-axis and sequential parasternal short-axis views were obtained to assess global and regional wall motion. Left ventricular (LV) dimensions at end-systole and end-diastole and fractional shortening (percent change in LV diameter normalized to end-diastole) were measured from the parasternal long-axis view using linear measurements of the LV at the level of the mitral leaflet tips during diastole. LV ejection fraction (EF), LV fractional shortening (FS), and LV mass are measured by using standard formulas for the evaluation of LV systolic function.^16^

### Histology

Hearts and lungs were fixed in cold 4% paraformaldehyde overnight at 4°C while shaking. The tissues were then washed multiple times with PBS at room temperature to remove paraformaldehyde. Paraffin sections (5-μm thickness) were stained with Hematoxylin/Eosin (H&E), reticulin, and Masson’s trichrome using reagents and kits from Sigma (Sigma, St. Louis, MO) following standard protocols. Images were taken using an Olympus IX70 microscope or Nikon eclipse 90i microscope.

Diagnosis of microvasculopathy was done by light microscopy. In microvessels (diameter 10 to 20μm), both the endothelial layer and the wall (medial) layer were identified. The endothelial layer was defined as the monocell layer at the inner part of the blood vessel wall. Endothelial cells were graded as thickened if the diameter of the cell layer was at least as thick as the endothelial cell cores. The wall layer (media) was defined as the polycell layer adjacent to the endothelium. Stenotic wall thickening was classified if the ratio of luminal radius to wall thickness was <1.^17^

### Stem cell transplantation assays

WT (CD45.1) recipient mice were irradiated with two doses of 540 rad 3h apart and then received 1 ×10^6^ unfractionated marrow cells from Tie2FF1 or Tie2-Cre control (CD45.2) donor mice by standard intravenous tail vein injection using a 27G insulin syringe. Peripheral blood was obtained every 4 weeks after transplantation and CD45.2 percentage chimerism and complete blood counts were measured.

### Isolation and culture of murine lung ECs

Primary murine lung EC (CD45^−^CD31^+^) isolation was performed as we previously described.^18, 19^ Briefly, mice were euthanized and the chest was immediately opened through a midline sternotomy. The left ventricle was identified, and the ventricular cavity was entered through the apex with a 27-gauge needle. The right atrium was identified and an incision was made in the free wall to exsanguinate the animal and to allow the excess perfusate to exit the vascular space. The animal was perfused with 30 ml of cold PBS. The lung tissue was collected and minced finely with scissors. The tissue fragments were digested in DMEM medium containing 1 mg/mL Collagenase D (Roche, Switzerland), 1 mg/mL Collagenase/Dispase (Roche) and 25 U/mL DNase (Sigma) at 37°C for 2 hr with shaking, after which the suspension was homogenized by triturating. The homogenate was filtered through a 70μm nylon mesh (BD Biosciences, San Jose, CA) and pelleted by centrifugation (400g for 5 min). Cells were first depleted for CD45^+^ cells (Miltenyi Biotec, SanDiego, CA) and then positively selected for CD31^+^ cells (Miltenyi Biotec) using magnetically labeled microbeads according to the manufacturer’s protocol. Isolated ECs (CD45^−^CD31^+^) were cultured on 1% gelatin coated plates in complete EC medium as we previously described,^18^ with no medium change for the first 72 hrs to allow EC attachment followed by medium change every 2-3 days. Cells were re-selected for CD31^+^ cells when they reach >70-80% confluence (usually after 3-4 days of culture).

### Transcriptome analysis of cardiac ECs using RNA sequencing

Cardiac ECs (CD45^−^CD31^+^) were isolated from Tie2FF1 or Tie2-Cre control mice by tissue digestion and magnetic bead isolation as described above. Total ribonucleic acid (RNA) was extracted using the RNeasy mini kit (Qiagen, Hilden, Germany). RNA integrity and quantitation were assessed using the RNA Nano 6000 Assay Kit of the Bioanalyzer 2100 system (Agilent Technologies, CA, USA). For each sample, 400ng of RNA was used to generate sequencing libraries using NEBNext® Ultra™ RNA Library Prep Kit for Illumina® (New England BioLabs, MA, USA) following manufacturer’s recommendations. The clustering of the index-coded samples was performed on a cBot Cluster Generation System using PE Cluster Kit cBot-HS (Illumina) according to the manufacturer’s instructions. After cluster generation, the library preparations were sequenced on an Illumina platform. Index of the reference genome was built using hisat2 2.1.0 and paired-end clean reads were aligned to the reference genome using HISAT2. HTSeq v0.6.1 was used to count the reads numbers mapped to each gene. Differential expression analysis between wild-type control ECs (from Tie2-Cre mice, n=3) and JAK2V617F-mutant ECs (from Tie2FF1 mice, n=3) was performed using the DESeq R package (1.18.0). The resulting *P*-values were adjusted using the Benjamini and Hochberg’s approach for controlling the false discovery rate. Genes with an adjusted *P*-value < 0.05 found by DESeq were assigned as differentially expressed. Gene Ontology (GO) (http://www.geneontology.org/) and KEGG pathways (http://www.genome.jp/kegg/) enrichment analysis of differentially expressed genes was implemented by the ClusterProfiler R package. GO terms and KEGG pathways with corrected P-value less than 0.05 were considered significantly enriched.

### Shear stress application

Primary murine lung ECs (passage 3-6) were seeded on 1% gelatin coated 6-well culture plates and allowed to grow to confluence. Unidirectional pulsatile shear stress at 60 dyne/cm^2^ was applied to confluent EC monolayers at 37 °C for 1 hr, using a programmable cone and plate shearing device.^20, 21^ The 60 dyne/cm^2^ shear stress was selected not only because it mimics the hemodynamic stress condition of mouse aortic arch blood flow,^22^ but also because it is compatible with our *in vitro* cell culture. Following shear exposure, EC monolayer was washed with PBS and collected for further testing.

### Gene expression analysis by real-time quantitative polymerase chain reaction

Total RNA was isolated using RNeasy Mini kit (Qiagen) following the manufacturer’s protocol. The TaqMan® Gene Expression Assay (Thermo Fisher Scientific, Waltham, MA) was used for real-time quantitative polymerase chain reaction (qPCR) to verify differential expression of Kruppel-like factors 2 (KLF2) and 4 (KLF4), thrombomodulin (TM), and eNOS on an Applied Biosystems 7300 Real Time PCR System (ThermoFisher Scientific). Values obtained were normalized to the endogenous Actin beta (Actb) gene expression and relative fold changes compared to control samples was calculated by the 2^ΔΔCT^ method. All assays were performed in triplicate.

### Flow cytometry analyses of blood and tissue samples

Expression of EC cell adhesion molecules PECAM-1 (platelet endothelial cell adhesion molecule, or CD31) and E-selectin was measured by flow cytometry using anti-mouse PECAM-1 (Clone 390, BD Biosciences) and anti-mouse E-selectin (Clone 10E9.6, BD Biosciences) antibodies and analyzed on a FACSCalibur (Becton Dickinson) cytometer.

### Isolation of murine HSPCs

12–18-week old mice were euthanized and the femurs and tibias removed. A 25-gauge needle was used to flush the marrow with PBS+2%FBS. Cells were triturated and filtered through 70 μM nylon mesh (BD Biosciences) to obtain a single cell suspension. For depletion of mature hematopoietic cells, the Lineage Cell Depletion Kit (Miltenyi Biotec) was used. The lineage (CD5, CD45R, CD11b, Ter119, and GR-1) negative cells were collected and then positively selected for CD117^+^ (cKit^+^) cells using CD117 microbead (Miltenyi Biotec) to yield Lineage^−^ cKit^+^ (Lin^−^cKit^+^) HSPCs.

### EC-HSPC co-culture in a mixed vascular environment in vitro

Three days prior to co-culture with HSPCs, 1×10^4^ primary murine lung ECs (passage 3-6) mixed from 0-100% JAK2V617F-mutant ECs (i.e. with wild-type: mutant EC ratio at 100:0, 90:10, 50:50, 0:100) were seeded into 1% gelatin coated 96-well plate in complete EC medium. One day prior to the co-culture experiment, Lineage^−^cKit^+^ (Lin^−^cKit^+^) HSPCs were isolated from Tie2-Cre control (wild-type) or Tie2FF1 (JAK2V617F) and cultured overnight in StemSpan serum-free expansion medium (SFEM) (Stem Cell Technologies) containing 100 ng/mL recombinant mouse SCF, 6 ng/mL recombinant mouse IL3 and 10 ng/mL recombinant human IL-6 (all from Stem Cell Technologies). On Day 0, 2,500 Lin^−^cKit^+^ cells were seeded onto the EC monolayer in 0.1mL SFEM with cytokines. 0.1 mL fresh SFEM with cytokines was added on Day 3 and cells were counted on Day 5.

### Statistical analysis

Statistical significance of differences in experiments with two groups and only one variable was assessed by unpaired Student’s t tests using Excel software (Microsoft). A p value of less than 0.05 was considered significant. For all bar graphs, data are presented as mean ± standard error of the mean (SEM). All experiments were conducted and confirmed in at least two replicates.

## RESULTS

### Development of spontaneous heart failure with increased risk of sudden death in the JAK2V617F-positive Tie2FF1 mice

Previously, to study the effects of JAK2V617F-bearing vascular microenvironment on MPN stem cell function, we crossed mice that bear a Cre-inducible human JAK2V617F gene (FF1)^14^ with Tie2-Cre mice^15^ to express JAK2V617F specifically in all hematopoietic cells (including HSPCs) and ECs (Tie2^+/−^FF1^+/−^, or Tie2FF1),^23^ so as to model the human diseases in which both the hematopoietic stem cells and ECs harbor the mutation. As we have reported, Tie2FF1 mice develop a myeloproliferative phenotype with leukocytosis, thrombocytosis, significant splenomegaly, and greatly increased numbers of hematopoietic stem cells by 8 wk of age.^23–25^

We noticed that there was an increased incidence of sudden death during performance of minor procedures (e.g. submandibular bleeding) in Tie2FF1 mice, especially after 20 wk of age (Figure 1A). This observation prompted us to examine the cardiac function of these mice using transthoracic echocardiography. At 20 wk of age, Tie2FF1 mice Tie2FF1 mice displayed a phenotype of dilated cardiomyopathy (DCM) with significant decreases in LV EF (47% *vs.* 69%, *P* = 0.004) and FS (23% *vs.* 38%, *P* = 0.006), and increases in LV end-diastolic volume (78 μl *vs.* 56 μl, *P* = 0.003) and end-systolic volume (41 μl *vs.* 18 μl, *P* = 0.0008) compared to age-matched Tie2-Cre controls. There was no echocardiographic evidence for regional wall motion abnormalities including segmental akinesis, dyskinesis, thinned or scarred wall, suggesting there was no evidence of transmural myocardial infarction. The cardiac dysfunction in Tie2FF1 mice persisted at 30wk of age. Since JAK2 is important for normal heart function,^26, 27^ we examined the mice at a younger age to be certain that the cardiovascular dysfunction we observed in Tie2FF1 mice was not due to developmental abnormalities. No significant differences in echocardiographic parameters were observed between Tie2FF1 and control mice at 10wk of age, suggesting that the DCM we observed in Tie2FF1 mice was not due to developmental abnormalities. (Figure 1B-C) In addition, the cardiomyopathy was not due to anemia (which is a cause of high output heart failure), as there was no significant differences in red blood cell count or hemoglobin level. (Figure 1D)

**Figure 1.**
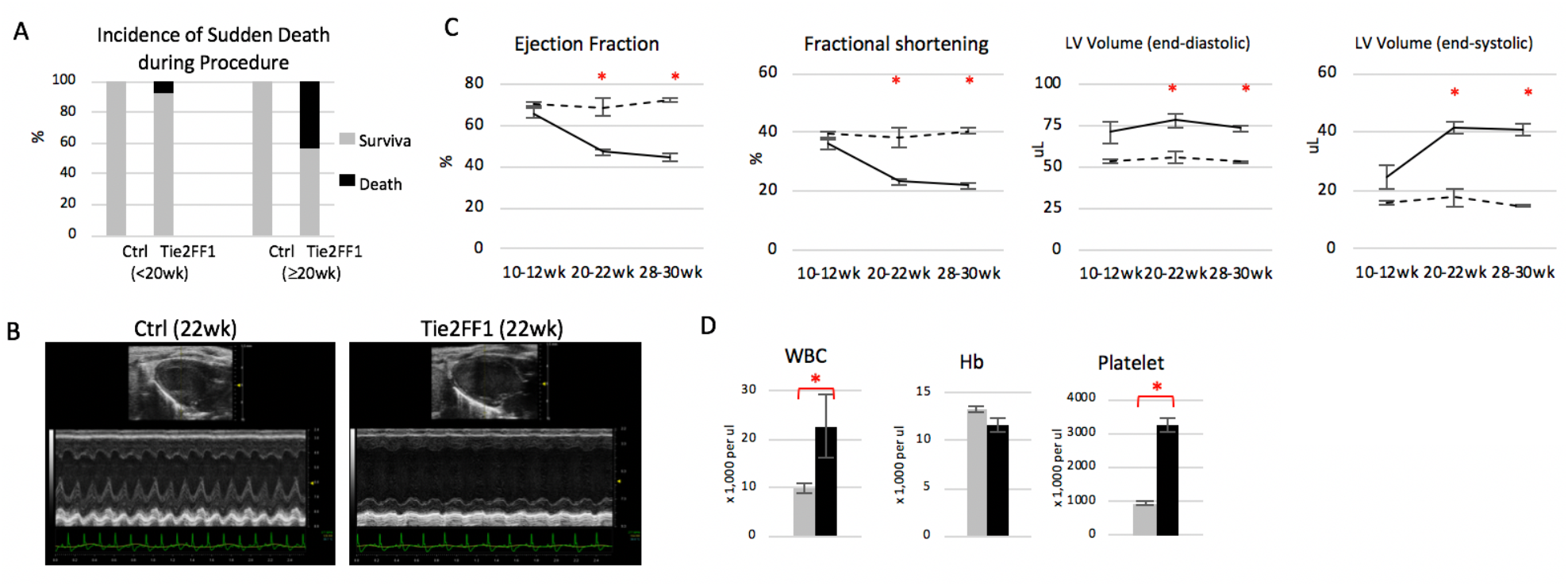
Development of spontaneous congestive heart failure in Tie2FF1 mice. (**A**) Increased incidence of sudden death in Tie2FF1 mice after 20wk of age. (**B-C**) Representative left ventricular echocardiographic tracings (B) and measurement of ejection fraction, fractional shortening, and left ventricular (LV) volume in Tie2 ctrl (dotted line) and Tie2FF1 (black line) mice (C). (n=3-4 mice in each group) (**D**) Blood counts of 20-22wk old Tie2-cre control (grey) and Tie2FF1 (black) mice (n=5-6 in each group). * P < 0.05

### JAK2V617F-positive Tie2FF1 mice have a prothrombotic and vasculopathy phenotype

Pathological evaluation confirmed the diagnosis of DCM with increased heart size and heart weight-to-tibia length ratio in Tie2/FF1 mice compared to controls. (Figure 2A) However, no significant atherosclerotic lesions (Figure 2B) or myocardium infarctions were detected in Tie2FF1 mice, which are consistent with echocardiographic findings.

**Figure 2.**
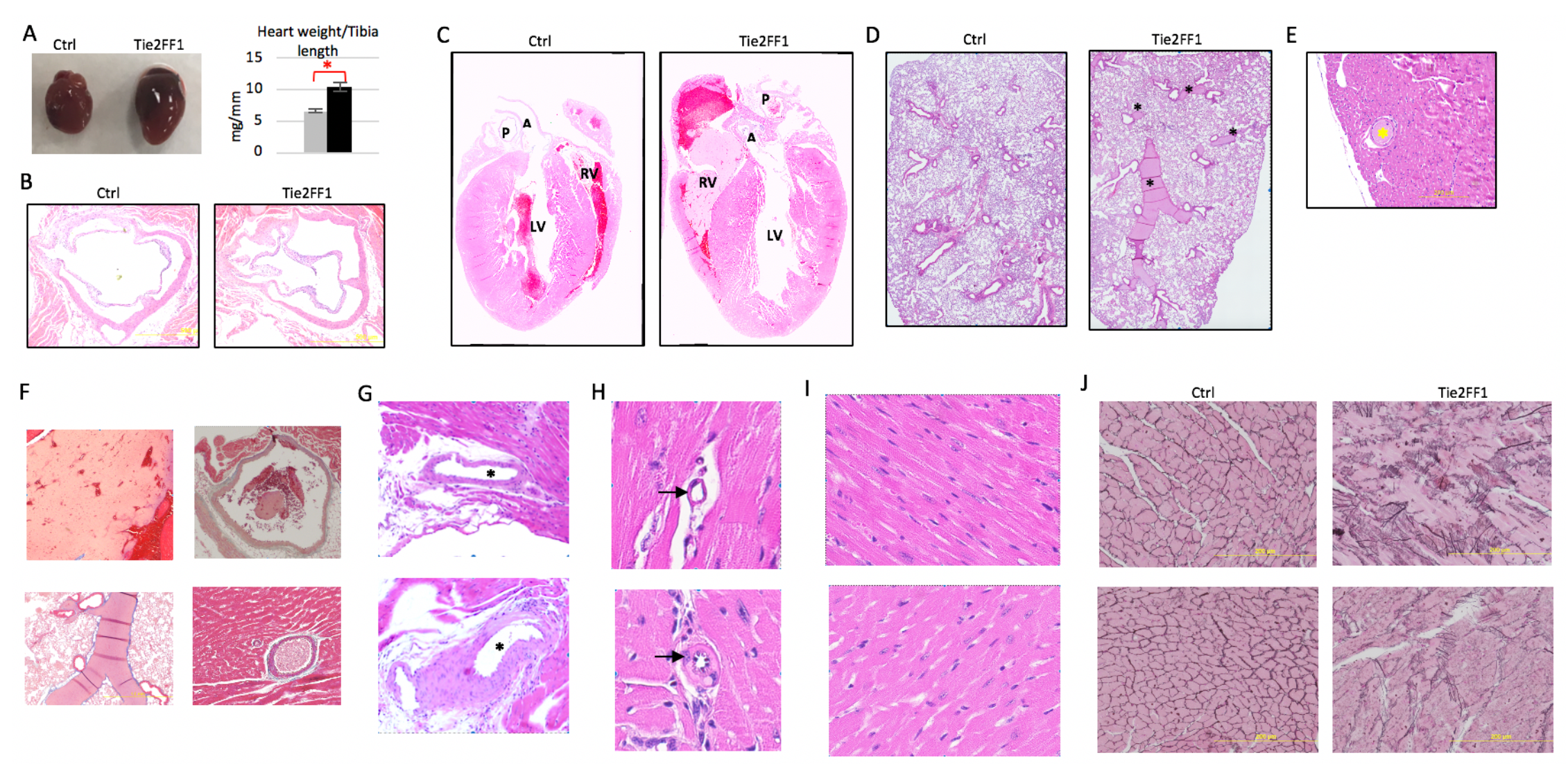
The JAK2V617F-positive Tie2FF1 mice have a prothrombotic and vasculopathy phenotype. (**A**) Representative image of heart (left) and heart weight adjusted by tibia length (right) of 20-30wk old Tie2 ctrl (grey) and Tie2FF1 (black) mice (n=4-5 in each group). (**B**) Representative H&E-stained aortic root area of Tie2-Cre ctrl and Tie2FF1 mice (magnification 4x). No significant atherosclerotic lesion was detected. (**C**) Representative H&E staining of longitudinal sections of Tie2-cre control and Tie2FF1 mice. Note the presence of thrombus in right ventricle and main pulmonary artery of the Tie2FF1 mice. LV: left ventricle; RV: right ventricle; A: aortic root; P: main pulmonary artery (magnification 10x). (**D**) Representative H&E staining of lung sections from Tie2-cre control and Tie2FF1 mice. Note the presence of thrombus (**\***) in segment pulmonary arteries of the Tie2FF1 mice (magnification 10x). (**E**) Representative H&E staining of coronary arteriole thrombus from the Tie2FF1 mice. (magnification 10x) (**F**) Representative masson’s trichrome staining demonstrates typical fibrin clot in right ventricle (upper left, 4x), pulmonary artery (upper right, 4x), segment pulmonary arteries (bottom left, 4x), and coronary arteriole (bottom right, 40x) of Tie2FF1 mice. (**G**) Representative H&E staining of epicardial coronary arteries (stars) in Tie2-cre control (top) and Tie2FF1 (bottom) mice. Note significant vascular wall thickening of the epicardial coronary artery in Tie2FF1 mice. (magnification 4x) (**H**) Representative H&E staining of intramyocardial coronary arterioles (arrows) in Tie2-cre control (top) and Tie2FF1 (bottom) mice. Note stenotic lumen narrowing of the arteriole in Tie2FF1 mice. (magnification 40x) (**I**) Representative H&E-stained cardiac sections showing enlarged cardiomyocytes in Tie2FF1 mice (bottom) compared to control mice (top) (magnification 40x). (**J**) Representative reticulin staining of heart sections from control (left) and Tie2FF1 (right) mice are shown. (magnification 20x)

We conducted several studies to determine the cause(s) for the sudden death and DCM phenotypes we observed in Tie2FF1 mice. First, we determined whether the JAK2V617F-positive Tie2FF1 mice developed spontaneous pulmonary thromboembolism, which is a common cause of sudden death.^28^ We found increased thrombosis in the right ventricle and both main and segment pulmonary arteries in Tie2FF1 mice, but not in age-matched Tie2-cre control mice. (Figure 2C-D and 2F) In addition, although we did not observe thrombosis in the main (epicardial) coronary arteries, we found thrombosis in scattered coronary arterioles (microvessels) in some of the Tie2FF1 mice examined. (Figure 2E and 2F) Such microvascular thrombosis in the coronary circulation can contribute to the development of dilated cardiomyopathy and heart failure.^29, 30^ Taken together, these results indicate that the JAK2V617F-positive Tie2FF1 mice have a pro-thrombotic phenotype affecting both the arterial and venous circulations.

We next examined the coronary vasculature of both Tie2FF1 and age-matched Tie2-cre control mice by light microscopy. Compared to control mice, Tie2FF1 mice demonstrated significant vascular smooth muscle cell proliferation and media wall thickening of their epicardial coronary arteries. (Figure 2G) In addition, there was significant stenotic lumen narrowing (i.e. ratio of luminal radius to wall thickness <1) involving scattered intramyocardial coronary arterioles in Tie2FF1 mice. (Figure 2H) These findings indicate that there are both macro- and micro-vasculopathy in Tie2FF1 mice.

Microscopic examinations of heart sections revealed enlarged cardiomyocytes and increased reticulin fibers in Tie2FF1 mice compared to control mice, (Figure 2I-J) reflecting the cardiac remodeling effects of both microvascular thrombosis and vasculopathy we observed in these mice. Such pathological cardiac remodeling can further impair cardiac function of Tie2FF1 mice.

### JAK2V617F-mutant ECs are required to develop the cardiovascular complications in Tie2FF1 mice

JAK2V617F is also one of the common mutations associated with CHIP.^2–6^ Over the past two years, in studies designed to better understand heart disease in individuals with CHIP, it has been reported that cardiovascular disease (e.g. atherosclerosis, heart failure) can develop in murine models with hematologic mutations (e.g. TET2, JAK2V617F) when the mice are challenged with additional risk factors (e.g. high-fat/high-cholesterol diet,^6, 31^ surgical ligation/constriction of the coronary artery or aorta,^32, 33^ or hyperlipidemia genetic background via knocking-out of low-density lipoprotein receptor^34^). In contrast, our Tie2FF1 mice, in which the JAK2V617F mutation is expressed in both blood cells and vascular ECs, develop spontaneous heart failure with increased risk of sudden death when fed on a regular chow diet. This difference suggests that mutant ECs can accelerate the cardiovascular pathology in this JAK2V617F-positive MPN murine model. To begin to address this hypothesis, we generated a chimeric murine model with JAK2V617F-mutant blood cells and wild-type vascular ECs by transplanting Tie2FF1 marrow cells into wild-type recipients. The transplantation of wild-type marrow cells into wild-type recipients served as a control. (Figure 3A) Recipients of Tie2FF1 marrow developed significant neutrophilia, monocytosis, and thrombocytosis after transplantation (Figure 3B), a peripheral blood phenotype virtually identical to primary Tie2FF1 mice (data not shown). However, echocardiographic evaluation did not reveal any differences in cardiac functional parameters between recipients of Tie2FF1 marrow and recipients of wild-type marrow during 8-month follow up after transplantation. (Figure 3C) Histology examination at the time of harvest did not reveal evidence of spontaneous thrombosis or coronary vasculopathy (data not shown). These results suggest that JAK2V617F-mutant blood cells alone are not sufficient to generate the spontaneous cardiovascular disease phenotype we observed in Tie2FF1 mice; JAK2V617F-mutant ECs are required.

**Figure 3.**
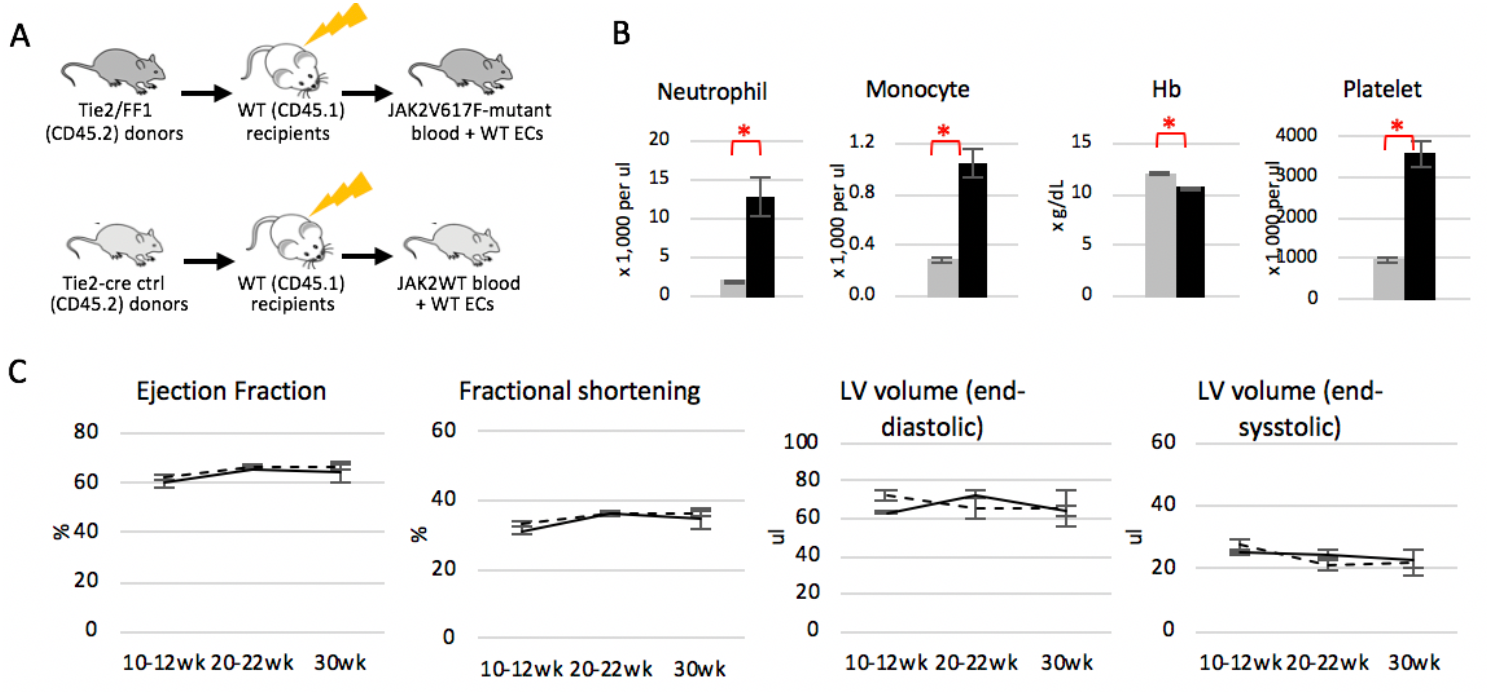
Normal heart function in mice with JAK2V617F-mutant blood cells and wild-type vascular ECs. (**A**) Experimental scheme to generate a chimeric murine model with JAK2V617F-mutant blood cells and wild-type vascular endothelium. (**B**) Blood counts in recipient mice of either Tie2-cre control (grey) or Tie2FF1 (black) marrow cells 22wk after transplantation. (**C**) Serial measurements of ejection fraction, fractional shortening, and left ventricular (LV) volumes in recipients of Tie2-cre control marrow (dotted line) and recipients of Tie2FF1 marrow (black line). (n=5 in each group) * *P* < 0.05

### The JAK2V617F mutation alters endothelial function

To understand the roles of JAK2V617F-mutant ECs in the development of cardiovascular complications in Tie2FF1 mice, transcriptomic profiles of wild-type and JAK2V617F-mutant cardiac ECs were analyzed using RNA sequencing. 234 genes from a total of 20,698 were differentially expressed (219 up- and 15 down-regulated) in JAK2V617F-mutant ECs compared to wild-type ECs. Dysregulation of leukocyte migration/chemotaxis, platelet activation, cell adhesion molecules, and blood coagulation pathways are highly enriched in JAK2V617F-mutant ECs, consistent with the prothrombotic phenotype we have observed in the Tie2FF1 mice. In addition, pathways involved in receptor-extracellular matrix interaction and leukocyte transendothelial migration/activation were also dysregulated in JAK2V617F-mutant ECs, which can contribute to the vasculopathy phenotype in these mice. (Figure 4A-B) These findings supported recent transcriptomic profiling of JAK2V617F-mutant human ECs derived from induced pluripotent stem cell lines from a MPN patient.^35^

**Figure 4.**
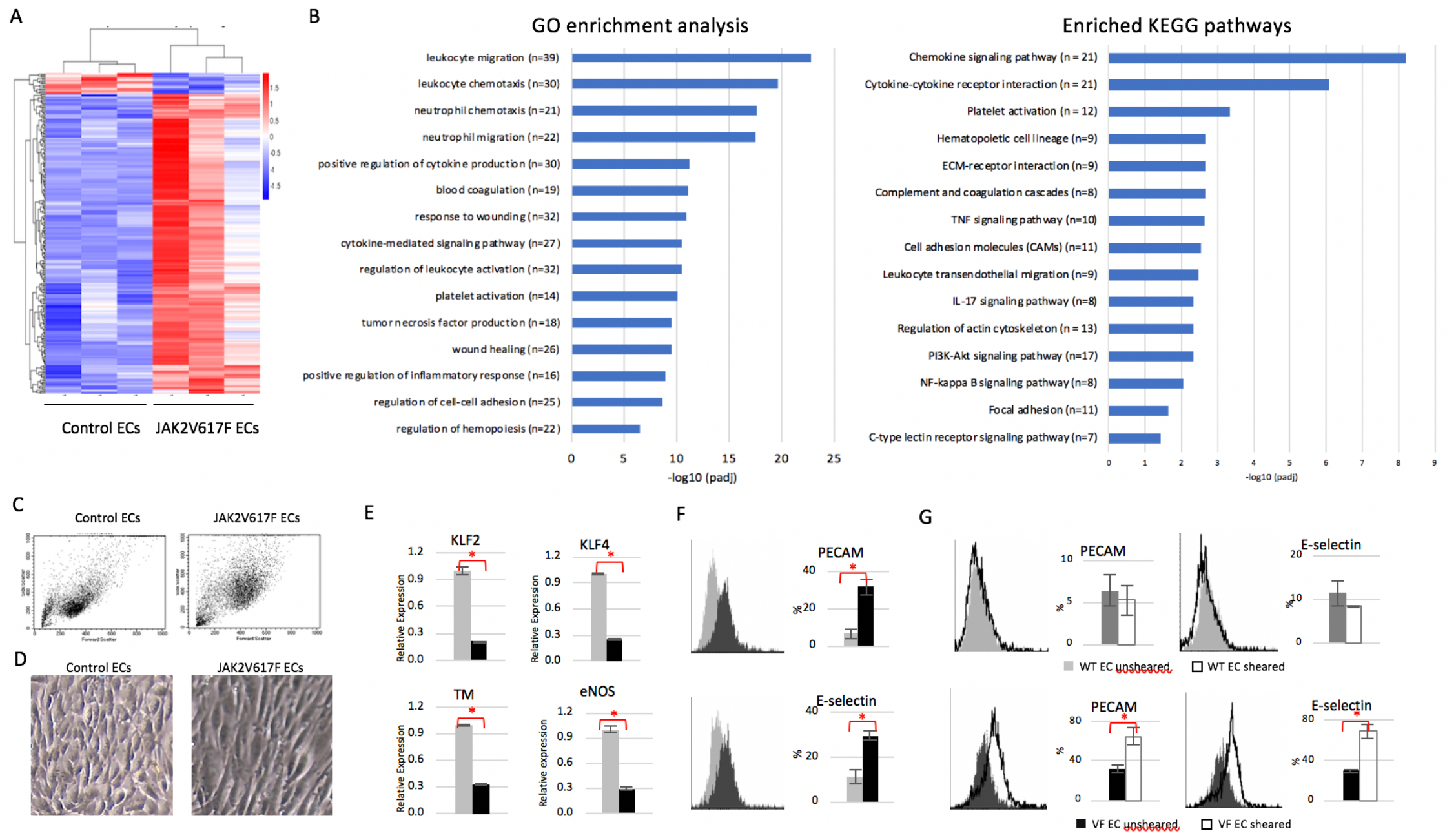
The JAK2V617F mutation alters endothelial function. (**A**) Non-supervised heat map and hierarchical clustering of 234 differentially expressed genes between wild-type control and JAK2V617F-mutant cardiac ECs with an adjusted *P*-value < 0.05. (**B**) Differentially enriched GO terms and KEGG pathways in JAK2V617F-mutant cardiac ECs compared to wild-type control ECs. P values are plotted as the negative of their logarithm. (**C**) Representative flow cytometry dot plots of wild-type and JAK2V617F-mutant ECs. Note JAK2V617F ECs have higher forward scatter (i.e. increased cell size) and side scatter (i.e. increased granularity) compared to wild-type ECs. (**D**) Representative bright field images of wild-type and JAK2V617F ECs (magnification: 10x). (**E**) Gene expression levels in wild-type (grey) and JAK2V617F (black) murine lung ECs were measured using realtime qPCR. Gene expression in VF ECs is shown as the fold change compared to WT EC expression which was set as ‘1’. (**F**) Representative flow cytometry histogram plots (left) and quantitative analysis (right) of PECAM (top panel) and E-selectin (bottom panel) expression in un-sheared wild-type (grey) and JAK2V617F (black) ECs. (**G**) Representative flow cytometry histogram plots and quantitative analysis of PECAM and E-selectin expression in unsheared vs. sheared wild-type (top panel) and JAK2V617F (bottom panel) ECs. * *P* < 0.05

We further studied how the JAK2V617F mutation alters endothelial function using primary murine lung ECs, in part because of their high proliferative activity^36^ and close resemblance to cardiac ECs in transcriptional profiling studies.^37^ First, we examined EC morphology in culture. Endothelial cell area correlates with cell stiffness and predicts cellular traction stress, which are involved in cell adhesion and migration.^38, 39^ Both microscopic examination and flow cytometric analysis revealed that JAK2V617F-mutant ECs are larger than wild-type ECs when seeded at similar cell density. (Figure 4C-D)

Next, we examined the gene expression levels of several key endothelial cell functional regulators. Kruppel-like factors 2 (KLF2) and 4 (KLF4) are important regulators of vascular function and confer an anti-atherogenic and anti-thrombotic endothelial phenotype.^40, 41^ We measured the expression levels of KLF2 and KLF4 in freshly isolated lung ECs from Tie2FF1 and Tie2-Cre control mice. Both KLF2 (0.21-fold, *P*<0.001) and KLF4 (0.25-fold, *P*<0.001) mRNA levels were significantly reduced in JAK2V617F-mutant ECs compared to wild-type ECs. (Figure 4E)

Thrombomodulin (TM) and endothelial nitric oxide synthase (eNOS) are two downstream targets of KLF2/4 signaling.^41^ Thrombomodulin is a vascular endothelial glycoprotein receptor that binds and neutralizes the prothrombotic actions of thrombin, and activates the natural anticoagulant protein C.^42^ Disturbances of the thrombomodulin–protein C antithrombotic mechanism are the most important known etiology for clinical disorders of venous thrombosis.^43^ eNOS plays an essential role in the regulation of endothelial function through the generation of nitric oxide, which inhibits platelet and leukocyte adhesion to the vessel wall and inhibits vascular smooth muscle cell proliferation.^44, 45^ Decreased NO as a consequence of impairment in eNOS is a hallmark of many cardiovascular diseases.^46^ Consistent with the pro-thrombotic and vasculopathy phenotype we have observed in Tie2FF1 mice, both TM (0.32-fold, P<0.001) and eNOS (0.30-fold, P<0.001) mRNA levels were significantly reduced in JAK2V617F-mutant ECs compared to wild-type ECs. (Figure 4E)

Last, we evaluated how the JAK2V617F mutation affects EC response to flow shear stress, which plays a critical role in endothelial function and atherothrombosis.^47^ Platelet-endothelial cell adhesion molecule (PECAM)^48, 49^ and endothelial cell selectin (E-selection)^50^ are two cell adhesion molecules involved in the regulation of thrombus formation and the pathogenesis of cardiovascular diseases. Quantitative flow cytometric analysis revealed that both PECAM (4.9-fold, *P*=0.038) and E-selectin (2.6-fold, *P*=0.038) were significantly up-regulated in JAK2V617F-mutant ECs compared to wild-type ECs. (Figure 4F) We then subjected cultured ECs to pulsatile unidirectional shear stress at 60 dyne/cm^2^, which mimic the hemodynamic condition of mouse aortic arch blood flow.^22^ We found that both PECAM (2.1-fold, *P*=0.075) and E-selectin (2.3-fold, *P*=0.027) were further up-regulated in sheared JAK2V617F-mutant ECs compared to un-sheared ECs. In contrast, their levels did not change in wild-type ECs. (Figure 4G) The pro-adhesive phenotype of JAK2V617F-mutant ECs we report here is supported by two recent studies using different experimental systems: JAK2V617F-mutant human ECs derived from induced pluripotent stem cell lines from MPN patient,^35^ and genetically engineered JAK2V617F-mutant human umbilical vein ECs.^51^

Taken together, these data indicate that the JAK2V617F mutation alters endothelial function, which could contribute to the cardiovascular complications observed in the Tie2FF1 mice.

### JAK2V617F-mutant ECs promote the expansion of JAK2V617F HSPCs in preference to wild-type HSPCs in a mixed vascular environment *ex vivo*

The JAK2V617F mutation is present in all blood cells and ECs of the Tie2FF1 mice, while the vascular endothelium in MPN patients is likely to be heterogeneous with co-existing normal and mutant ECs.^8–11, 13, 52^ To better mimic the human disease, we designed an *ex vivo* co-culture system with a mixed vascular microenvironment. We again utilized murine lung ECs, not only because of their high proliferative endothelial progenitor activity in lung microvasculature^36^ and a recent report that lung is an organ with considerable hematopoietic potential,^53^ but also because JAK2V617F-mutant murine lung ECs have successfully recapitulated the neoplastic hematopoiesis of human MPN disease (i.e. MPN stem cell expansion and disease relapse after marrow transplantation) *in vitro* in our previous studies.^19, 24, 25^ When wild-type or JAK2V617F-mutant Lin^−^cKit^+^ HSPCs were seeded on a monolayer of lung endothelium mixed from 0-100% JAK2V617F-mutant ECs, we found that the presence of as low as 10% mutant ECs in the mixed monolayer promoted the expansion of mutant HSPCs in preference to wild-type HSPCs. (Figure 5) This finding was not surprising to us, since as few as 2% mutant blood cells can produce >2-fold increase of heart disease in individuals with CHIP. Therefore, we reasoned, and confirmed that even small numbers of JAK2V617F-mutant ECs can mimic one of the MPN disease phenotypes, that is, mutant EC promotion of mutant HSPC proliferation.

**Figure 5.**
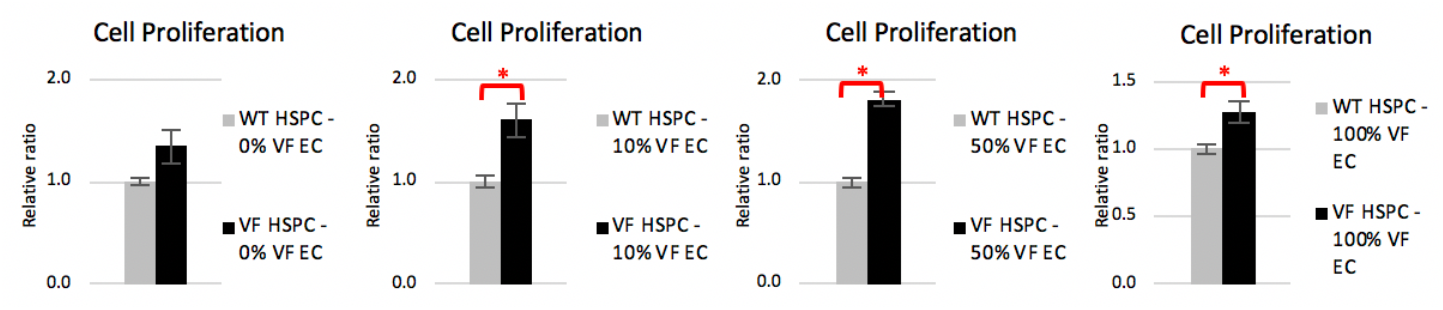
Lineage^neg^cKit^+^ HSPC cell proliferation *in vitro* in a mixed vascular environment where wild-type (WT) and JAK2V617F-mutant (VF) lung ECs are mixed at different ratios (with 0-100% mutant ECs). VF HSPC cell proliferation was shown as the relative ratio compared to WT HSPCs cultured under the same conditions. Data are from one of two independent experiments that gave similar results. * *P*<0.05

### Stem cell transplantation improves cardiac function in the JAK2V617F-positive Tie2FF1 mice

Stem cell transplantation (SCT) is a promising therapeutic strategy to repair impaired myocardium in patients with heart failure.^54–57^ SCT is also the only curative treatment for patients with MPNs.^58^ It is not known whether SCT can improve cardiac function in these patients who suffer from many cardiovascular complications during the course of their diseases.

We investigated whether SCT can improve cardiac function in Tie2FF1 mice by transplanting wild-type marrow cells into 20-26wk old Tie2FF1 mice with DCM. The transplantation of wild-type marrow cells into age-matched Tie2-cre mice served as a control. (Figure 6A) Cardiac dysfunction (i.e. decreases in LVEF and fractional shortening, and increases in left ventricular end-diastolic volume and end-systolic volume) gradually improved in Tie2FF1 recipient mice during 7-month follow up. (Figure 6B) In addition, there was no significant difference in heart weight between the two groups at 7-month post transplantation. (Figure 6C) At 7-month post transplantation, although there was still stenotic vasculopathy involving scattered coronary arterioles in Tie2FF1 recipient mice, no significant difference in cardiomyocyte size or reticulin fibers was detected between the two groups. (Figure 6D-F) Importantly, this beneficial effect of SCT was obtained despite MPN disease relapse as we previously reported in these mice.^25^ (Figure 6G-H)

**Figure 6.**
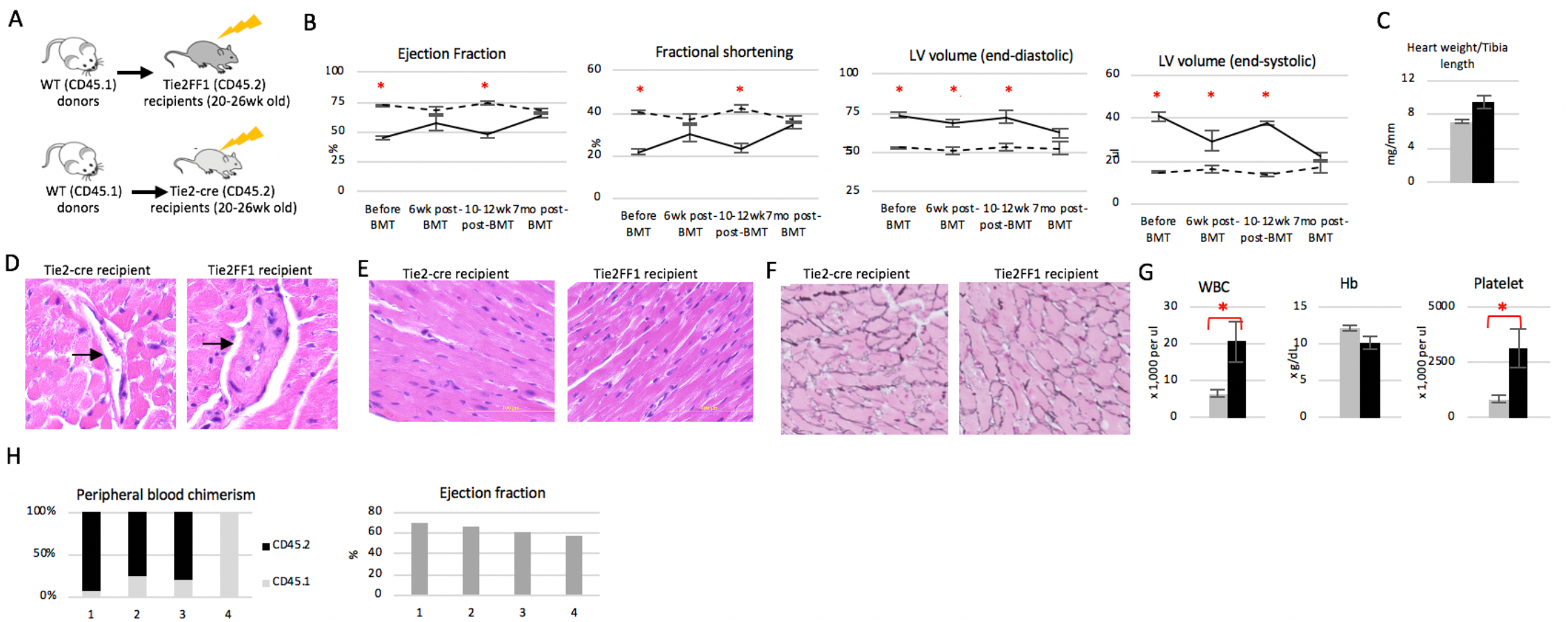
Stem cell transplantation improves cardiac function in the JAK2V617F-positive Tie2FF1 mice. (**A**) Experimental scheme of stem cell transplantation in heart failure 20-26wk old Tie2FF1 mice. (n=4 in each group) (**B**) Serial measurements of ejection fraction, fractional shortening, and left ventricular (LV) volumes in Tie2-cre recipients (dotted line) and Tie2FF1 recipients (black line). (**C**) Heart weight adjusted by tibia length of Tie2-cre recipients (grey) and Tie2FF1 recipients (black) 7-month after transplantation. (**D**) Representative H&E staining of coronary arterioles (arrows) in Tie2-cre control and Tie2FF1 recipient mice. (magnification 40x) (**E**) Representative H&E-stained cardiac sections corresponding to the left ventricular free wall of hearts in Tie2-cre control and Tie2FF1 recipient mice. (magnification 40x) (**F-g**) Representative reticulin staining of heart sections from Tie2-cre control and Tie2FF1 recipient mice. (magnification 20x) (**G**) Blood counts in Tie2-cre (grey) and Tie2FF1 (black) recipient mice 7 months after transplantation. (**H**) Peripheral blood chimerism (left) and heart ejection fraction (right) in four Tie2FF1 recipient mice 7-month after transplantation. 3 of the 4 Tie2FF1 recipient mice had disease relapse though there was no difference in cardiac function between relapsed and non-relapsed Tie2FF1 recipients.

Taken together, stem cell transplantation (with wild-type donor marrow cells) can reverse adverse cardiac remodeling (i.e. attenuate fibrosis) and improve cardiac function in the Tie2FF1 mice with DCM.

## DISCUSSION

Up to 40% of patients with MPNs develop arterial or venous thrombosis during the course of their disease, with cardiovascular events being the leading cause of morbidity and mortality in these patients.^1^ Despite the substantial progress in our understanding of hemostasis and thrombosis, little is known of the mechanisms that contribute to the hypercoagulability and vascular complications in patients with MPNs, the actual cause of death in the majority of patients who succumb to their illness. Vascular endothelium plays important roles in the regulation of hemostasis and thrombosis. Endothelial cells carrying the JAK2V617F mutation can be detected in many patients with MPNs.^8–13, 52^ However, few studies have examined the role(s) of mutant ECs in the development of cardiovascular complications in patients with MPNs.

In this study, we utilized a transgenic murine model of MPN (Tie2FF1) in which the JAK2V617F mutation is expressed in both blood cells and vascular ECs, so as to model the human diseases in which both the blood cells and ECs harbor the mutation. We observed that Tie2FF1 mice develop spontaneous, age-related DCM with an increased risk of sudden death. (Figure 1) Pathologic evaluation revealed increased thrombosis in pulmonary arteries, right ventricle, and small coronary arterioles of the Tie2FF1 mice. There was also evidence of macro- and micro-vasculopathy in coronary vasculature of the Tie2FF1 mice manifested as vascular media wall thickening of the epicardial coronary arteries and stenotic lumen narrowing of the intramyocardial arterioles. (Figure 2) No significant myocardium infarctions were detected in Tie2FF1 mice. Therefore, massive pulmonary embolism and/or malignant ventricular arrythmia due to severely decreased LV systolic function are likely the causes of sudden death in these mice.

In contrast to other murine models of hematologic disorders (e.g. TET2- or JAK2V617F-induced CHIP), in which cardiovascular diseases (e.g. atherosclerosis, heart failure) only develop when the mice are challenged with additional risk factors,^6, 31–34^ our Tie2FF1 mice (with both JAK2V617F-mutant blood cells and JAK2V617F-mutant vascular ECs) develop spontaneous DCM when fed on regular chow diet. In contrast, a chimeric murine model with JAK2V617F-mutant blood cells and wild-type vascular ECs did not develop any sign of cardiac dysfunction during 8-month follow up despite the development of myeloproliferative phenotype. (Figure 3) These observations suggest that JAK2V617F-mutant ECs are required to develop/accelerate the cardiovascular pathology in this murine model of JAK2V617F-positive MPN. Previously, Shi and colleagues reported that a JAK2V167F-positive murine model of MPN (in which the mutation is expressed only in blood cells under the control of the *Vav-1* promoter) developed cardiomegaly and coronary artery thrombosis on histology examination (though no functional evaluation of heart function was provided).^59^ One possible explanation for the difference between our observation and Shi et al.’s study is the marked erythrocytosis present in Shi’s murine model, which is a primary determinant of blood viscosity and a contributor to vascular complications in patients with MPNs.^60^

To help explain the molecular mechanism(s) by which the JAK2V617F mutation alters endothelial function to promote cardiovascular complications in MPNs, we found that: (1) the key endothelial function regulators KLF2, KLF4, TM, and eNOS levels were significantly down regulated in JAK2V617F-mutant ECs compared to wild-type ECs; (2) the cell surface adhesion molecules PECAM and E-selectin were significantly upregulated in the mutant ECs; and (3) both PECAM and E-selectin levels were further upregulated in JAK2V617F-mutant ECs by flow shear stress, while their levels did not change in wild-type ECs. (Figure 4) Together with our previous findings that JAK2V617F-mutant ECs have increased cell proliferation, cell migration, and angiogenesis *in vitro* compared to wild-type ECs,^19, 24, 25^ our work demonstrates that the JAK2V617F mutation promotes a pro-proliferative, pro-adhesive, and pro-thrombotic endothelial phenotype that can play important roles in the development of cardiovascular complications in MPNs.

Both animal studies and clinical trials have reported that stem cell transplantation can improve heart function though the underlying mechanism remains unclear.^56, 57^ It has been shown that stem cells can exert paracrine effects by secreting cardio-protective factors, ^61, 62^ or stimulate local inflammatory response in injured heart to improve its function.^63^ In our heart failure Tie2FF1 mice, SCT with wild-type donor marrow cells reversed cardiac fibrosis and improved heart function gradually during 7 months follow up. Consistent with our previous report on JAK2V617F-mutant ECs in MPN disease relapse after SCT,^25^ 3 of the 4 Tie2FF1 recipient mice had disease relapse with mixed chimerism, leukocytosis, and thrombocytosis 7-month post transplantation. However, there was no difference in cardiac function between relapsed and non-relapsed Tie2FF1 recipient mice, suggesting the beneficial effect of SCT can be obtained despite MPN disease relapse. Pathology examination revealed that SCT reversed the left ventricle remodeling process in this JAK2V617F-positive heart failure MPN murine model. (Figure 6) Considering SCT is the only curative treatment for patients with MPNs and many of these patients were excluded from SCT due to poor cardiac function, further clinical and laboratory investigation is very much needed.

Though a common origin between blood and endothelium beyond the embryonic stage of development continues to be a subject of intense investigation, the available data is clear that JAK2V617F-mutant ECs are present in many patients with MPNs, albeit at variable frequency.^8–13^ ECs are also involved by the malignant process in many other hematologic malignancies.^65–67^ Vascular ECs in patients with MPNs are likely heterogeneous with both normal and JAK2V167F-mutant ECs.^9^ Since as low as 2% mutant blood cells (in that case, monocytes) can produce >2-fold increase of heart disease in individuals with CHIP,^64^ we reason that even small numbers of JAK2V617F-mutant ECs may affect the cardiovascular phenotype in MPN patients. Murine models of chimeric vascular endothelium (i.e. with both wild-type and JAK2V617F-mutant ECs) will be required to further test this hypothesis. Taken together with recent reports implicating vascular endothelium in the generation of thrombotic events in MPNs,^35, 51^ our findings suggest that the JAK2V617F mutation in ECs alters vascular endothelial function to promote cardiovascular complications including thrombosis, vasculopathy, cardiomyopathy in MPNs. Therefore, targeting the MPN vasculature represents a promising new therapeutic strategy in patients with MPNs.

## ACKNOWLEDGEMENTS

We thank Yan Ji from the Research Histology Core Laboratory at Stony Brook Medicine for assistance with our histological studies. We thank Eric Weber (Novogene Corporation Inc. CA, USA) for providing useful consultation on RNA sequencing data analysis. This research was supported by the National Heart, Lung, and Blood Institute grant NIH R01 HL134970 (H.Z.) and VA Merit Award BX003947 (H.Z.).

